# Linear dicentric chromosomes in bacterial natural isolates reveal common constraints for replicon fusion

**DOI:** 10.1101/2025.02.23.639760

**Authors:** Ram Sanath-Kumar, Arafat Rahman, Zhongqing Ren, Ian P. Reynolds, Lauren Augusta, Clay Fuqua, Alexandra J. Weisberg, Xindan Wang

## Abstract

Multipartite bacterial genome organization can confer advantages including coordinated gene regulation and faster genome replication but is challenging to maintain. *Agrobacterium tumefaciens* lineages often contain a circular chromosome (Ch1), a linear chromosome (Ch2), and multiple plasmids. We previously observed that in some stocks of the lab model strain C58, Ch1 and Ch2 were fused into a linear dicentric chromosome. Here we analyzed *Agrobacterium* natural isolates from the French Collection for Plant-Associated Bacteria (CFBP) and identified two strains with fused chromosomes. Chromosome conformation capture identified integration junctions that were different from the C58 fusion strain. Genome-wide DNA replication profiling showed both replication origins remain active. Transposon sequencing revealed that partitioning systems of both chromosome centromeres are essential. Importantly, the site-specific recombinases XerCD are required for the survival of the strains containing the fusion chromosome. Our findings show that replicon fusion occurs in natural environments and that balanced replication arm sizes and proper resolution systems enable the survival of such strains.

**Importance:** Most bacterial genomes are monopartite with a single, circular chromosome. But some species, like *Agrobacterium tumefaciens,* carry multiple chromosomes. Emergence of multipartite genomes is often related to adaptation to specific niches including pathogenesis or symbiosis. Multipartite genomes confer certain advantages, however, maintaining this complex structure can present significant challenges. We previously reported a laboratory-propagated lineage of *A. tumefaciens* strain C58 in which the circular and linear chromosomes fused to form a single dicentric chromosome. Here we discovered two environmental isolates of *A. tumefaciens* containing fused chromosomes derived from a different route, revealing the constraints and diversification of this process. We found that balanced replication arm sizes and the repurposing of multimer resolution systems enable the survival and stable maintenance of dicentric chromosomes. These findings help us better understand how multipartite genomes function across different bacterial species and the role of genomic plasticity in bacterial genetic diversification.

## Introduction

Most bacteria have a single chromosome. However, approximately 10% of bacteria contain other replicons in addition to the main chromosome (1). These multipartite genomes are common in bacteria involved in pathogenesis and symbiosis (2, 3). Most multipartite genomes are stably maintained, although there are rare examples of bacterial taxa that undergo frequent replicon fusion and splitting (*e.g. Rhizobium sp*. strain NGR234) (4). Evolutionary drivers for the emergence of these complex genomes include adaptation to new niches, as seen in the nitrogen-fixing symbiont *Sinorhizobium meliloti*, human pathogens including *Vibrio cholerae* and *Brucella melitensis*, and plant pathogens like *Agrobacterium tumefaciens* (5–10).

A specific subset of *A. tumefaciens* lineages have a complex genome consisting of a circular chromosome (Ch1), a linear chromosome (Ch2), and two or more large plasmids (pAt, pTi, and others) (9, 11). We previously found that surprisingly, particular stocks of the commonly used lab strain *A. tumefaciens* C58 have undergone fusion of Ch1 and Ch2 into a stable linear dicentric chromosome over the course of laboratory cultivation (12). Here, we have identified two environmental *A. tumefaciens* isolates each containing a different fused chromosome. We analyzed their integration sites and examined genetic requirements for their survival. Comparing these natural isolates with our previously identified C58 fusion strain provides insights into the plasticity of and requirements for maintaining fusion chromosomes.

## Results

### Two natural isolates of *A. tumefaciens* contain a fused chromosome

To explore whether chromosome fusion naturally occurs in environmental isolates, we performed long-read nanopore whole-genome sequencing for over 400 *Agrobacterium* strains, including several from the French Collection for Plant-Associated Bacteria (CIRM-CFBP). We identified two strains, CFBP_2407 and CFBP_2642, isolated from grapevine galls in two geographically separated grape growing localities in Hérault and Gironde departments in France, respectively, that each had a single large linear chromosome in place of binary replicons (13). This genomic configuration was reminiscent of, but different from, the chromosomal fusion in C58 (12). To find the closest relative of these strains with a binary chromosome structure, we constructed a core genome phylogeny made from 4,192 core genes extracted from publicly available genomes. In this phylogeny, both CFBP_2407 and CFBP_2642 are most closely related to each other, with the closest sister clade containing strains 47-2, IL15, and 1D1418 (**Fig. 1A**). The three strains of this sister clade are all predicted to have typical bipartite genomes based on their long-read assemblies. We chose strain 47-2, isolated from a grapevine gall in Israel, as a comparator for CFBP_2407 and CFBP_2642. Contig synteny analysis identified conserved regions and revealed a high degree of similarity (>99.3% average nucleotide identity) between 47-2 and the fusion chromosome strains (**Fig. 1B**).

**Figure 1.**
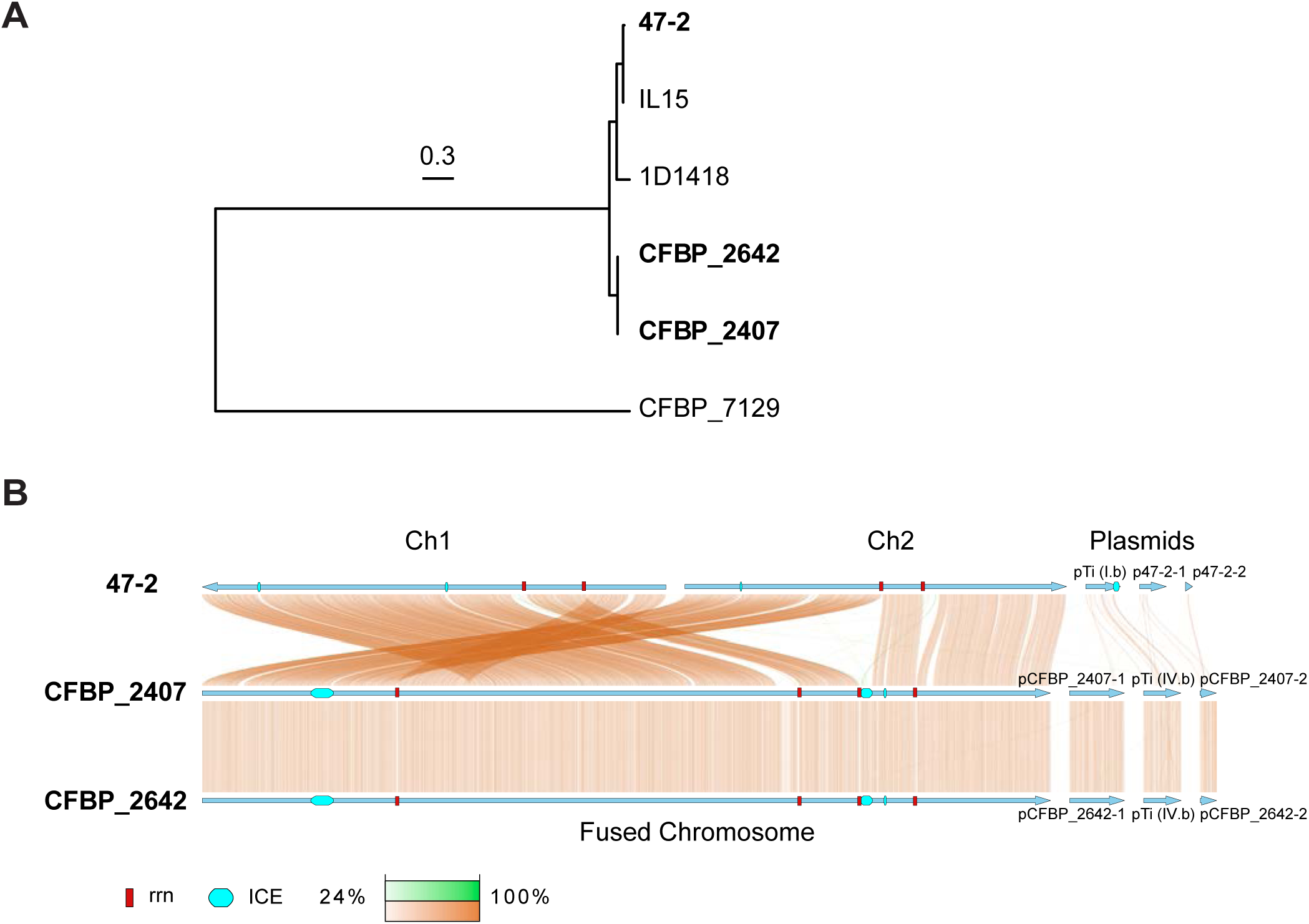
Phylogenetic analysis of natural *A. tumefaciens* isolates. **(A)** Maximum likelihood phylogeny based on 4,192 genes conserved in the 6 analyzed strains. The tree is midpoint-rooted. Branches with ultrafast bootstrap > 95% and SH-aLRT > 80% are black, branches with other support are gray. Branch scale is indicated by a bar. **(B)** Genome synteny between 47-2, CFBP_2407, and CFBP_2642. Blue arrows indicate individual replicons and their direction in the assembly. Colored bars linking replicons of different strains indicate regions of similarity. Copper colored links indicate the same orientation, green links indicate inversions. Darker colors indicate greater sequence similarity. Red boxes indicate the position of *rrn* regions. Cyan rounded rectangles indicate the position of integrative and conjugative elements (ICEs).

### Identifying the sites of chromosome fusion

To further investigate the genome architecture of these three strains, we performed genome-wide chromosome conformation capture (Hi-C) analysis. The 47-2 strain has a binary chromosome structure, as indicated by clear breakpoints between Ch1 (2,890 kb) and Ch2 (2,410 kb) (**Fig. 2A**). In contrast, both CFBP_2407 and CFBP_2642 exhibited a continuous genomic map of Hi-C data without any breakpoint, indicating the presence of a single, fused chromosome of ∼5,361 kb (**Figs. 2B** and **2C**). Consistent with our nanopore long-read assemblies, strains CFBP_2407 and CFBP_2642 lacked interactions at the termini of the chromosome, indicating a linear chromosome as opposed to the circular Ch1 in strain 47-2 (red circles on the maps of **Figs. 2B** and **2C**).

**Figure 2.**
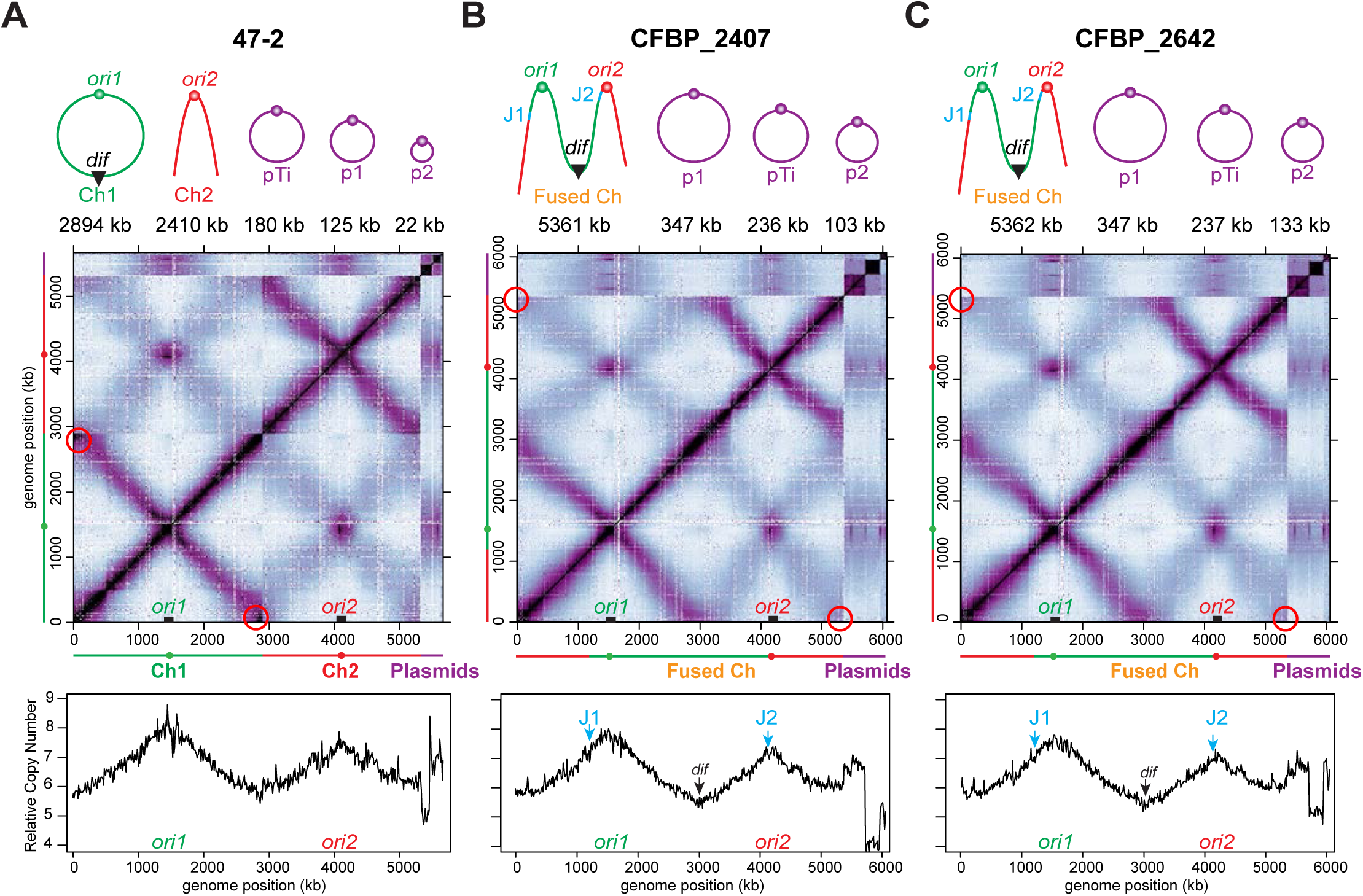
Natural *A. tumefaciens* isolates CFBP_2407 and CFBP_2642 exhibit a linear dicentric chromosomes. (A-C) CFBP_2407 and CFBP_2642 contain a fused chromosome, while 47-2 serves as a binary control. Top, schematics of genome composition of indicated strains. J1 and J2 mark the junctions of chromosome fusion. Middle, normalized Hi-C interaction maps of the respective genomes. Red circles indicate the terminus regions of the circular Ch1 **(A)** or the linear fusion chromosomes (**B, C**). Bottom, marker frequency analysis of exponentially growing cells. The y-axis depicts number of reads.

To identify the chromosome fusion sites, we mapped the Hi-C reads of CFBP_2407 and CFBP_2642 to the binary control strain 47-2. As expected, this comparison revealed a shuffled Hi-C map, indicating substantial genome reorganization in the fusion isolates compared with the binary control (**Fig. S1**). The shuffled Hi-C maps revealed two breakpoints at 2,156 kb on Ch1 and 1,161 kb on Ch2, where the fusion likely occurred (**Fig. 3**). These breakpoints are consistent with the synteny analysis derived from nanopore sequencing (**Fig. 1B**). Notably, both breakpoints were located within two ∼6-kb regions containing identical ribosomal RNA (*rrs*, *rrl*, and *rrf*) and tRNA genes (*trnI, trnA*, and *trnfM*) (**Fig. 3B-D**), indicating that the fusion event occurred by recombination of homologous regions on different chromosomes. Importantly, both CFBP_2407 and CFBP_2642 contain a 77-kb region that resembles an integrative and conjugative element (ICE) inserted immediately adjacent to the fusion junction (**Fig. 3D**). We hypothesize that integration of the ICE may have been a trigger for this fusion event. Notably, the fusion junctions in these natural isolates were different from those identified in the previously characterized lab strain C58, both in terms of location and gene content (12).

**Figure 3.**
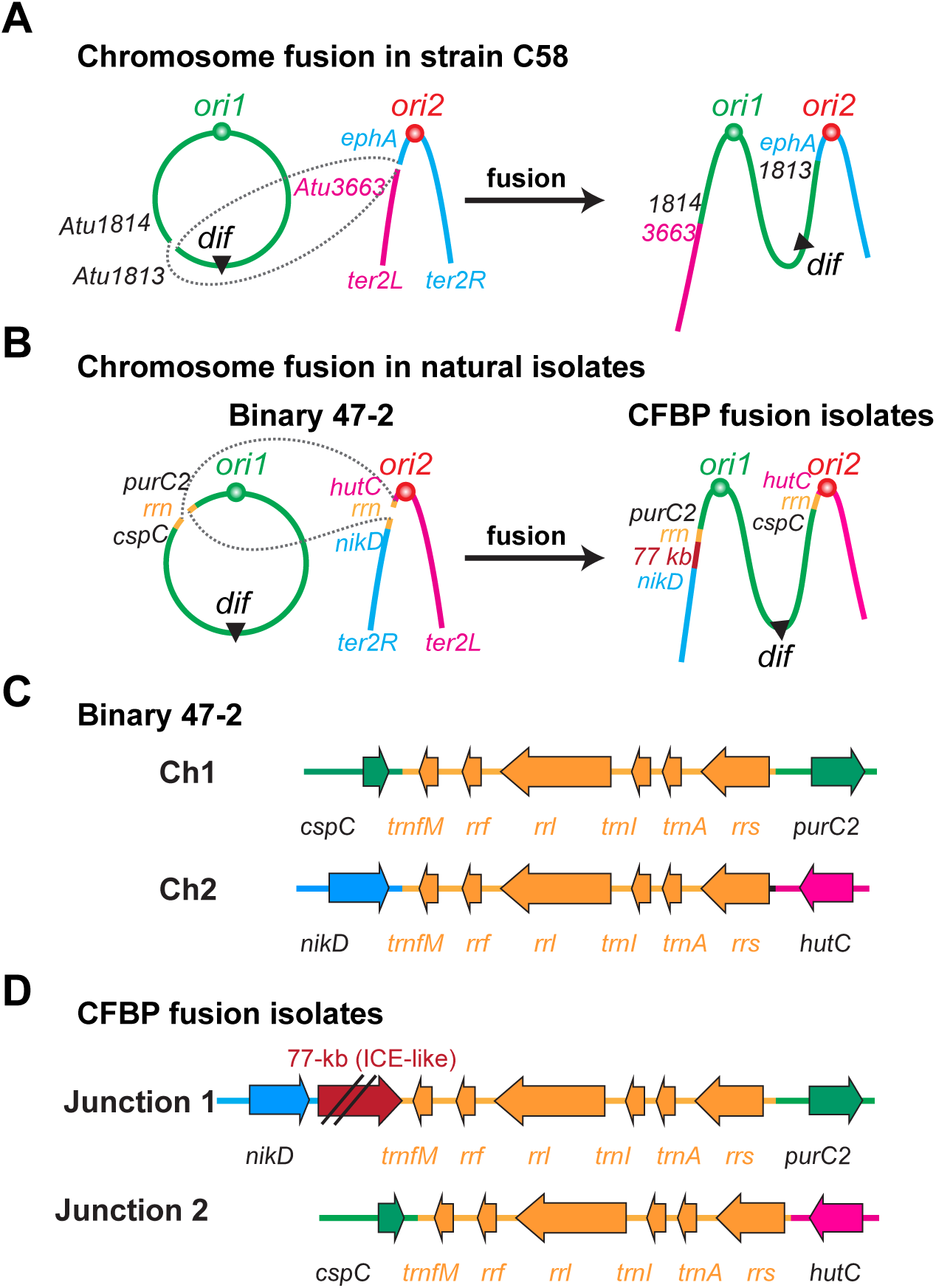
Sites of chromosome fusion in lab strains and natural isolates. **(A)** Schematic illustration of chromosome fusion events in C58 (12). **(B)** Schematic illustration of chromosome fusion events in the natural isolates of CFBP_2407 and CFBP_2642 compared with the binary control 47-2. CFBP fusion isolates have a 77-kb insertion (maroon) resembling an integrative conjugative element. **(C–D)** Genetic context of the fusion junctions in the binary and fusion strains. The 77-kb ICE region is not drawn to scale.

### The two replication origins on the fusion chromosomes remain active and are essential

A surprising finding from our analysis of the C58 fusion strain was that both origins of the fused chromosome independently initiate replication (12). To understand the replication profiles of these two fusion strains, we performed marker frequency analysis (MFA) using whole-genome sequencing. Both origins of replication (*ori1* and *ori2*) in the fusion chromosomes of CFBP_2407 and CFBP_2642 were firing at a similar frequency to those in the binary 47-2 strain (**Figs. 2A – C**, bottom), as signified by comparable origin copy numbers. All the strains had similar replication rates as denoted by the slope from each origin peak (**Figs. 2A – C**, bottom). To examine gene essentiality, we performed transposon sequencing (Tn-seq) of the two fusion strains and binary control strain 47-2. We found both centromeric partitioning systems – *parAB* for Ch1 and *repABC* for Ch2 – were essential in all three strains, regardless of chromosome configuration (**Figs. 4A – B**).Thus, the fusion chromosomes in the natural isolates remained the same replication and segregation programs as their binary counterparts.

**Figure 4.**
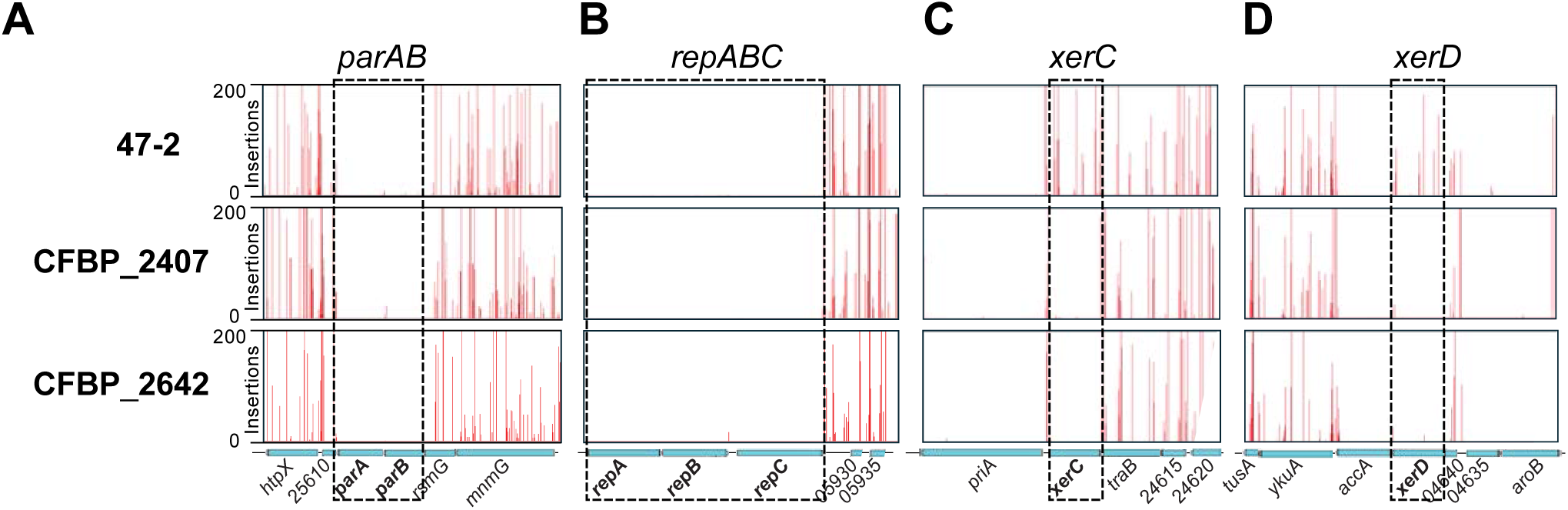
Gene essentiality in the three natural isolates. Transposon sequencing profiles at Ch1 partitioning system *parAB* **(A)**, Ch2 partitioning system *repABC* **(B)**, genes encodes for tyrosine recombinases *xerC* **(C)** and *xerD* **(D)**. The x-axis depicts genome position, and the y-axis represents the number of transposon insertions.

### XerC/D/*dif* recombination system is required for the viability of the fusion strains

Our Tn-seq results showed that the site-specific recombinases *xerCD* were essential in the fusion strains but not in the binary 47-2 strain (**Figs. 4C – D**). Importantly, sequence analysis showed that the *dif* site was located roughly midway between *ori1* and *ori2*, and at the convergence point of the FtsK Orienting Polar Sequences (KOPS) (**Fig. S2A**). This pattern of gene essentiality and KOPS orientation is consistent with our findings in the C58 fusion strain (12). The *ori1* and *ori2* being active and the essentiality of the two partitioning systems indicate that the two origins of the fusion chromosomes remain independent for replication and segregation. The independent *ori* function results in a situation where for every cell division, ∼50% of cells have the two replication origins of the fusion chromosome in opposite daughter cells (**Fig. S2B**). If this situation is unresolved, the affected cells will be impaired for chromosome segregation and cell division, which causes DNA breakage and cell death (**Fig. S2B**). The presence of the *dif* site at the convergence of KOPS sites near the closing septum enables FtsK to efficiently activate XerCD recombination at *dif* to resolve the partitioning issue, ensuring that each daughter cell receives a complete and properly partitioned genome (**Fig. S2B**).

## Discussion

In this study, we identified two natural *Agrobacterium* isolates with fused chromosomes analogous to our previous finding of a dicentric chromosome in the lab strain C58. The fact that two geographically distinct isolates have precisely the same change suggest that chromosome fusion occurred prior to the separation of these closely related isolates in the environment. Comparisons between the three fusion strains reveal several similarities of these fused chromosomes (**Figs. 3A – B**): 1) the recombination that resulted in the fusions occurred between highly similar sequences on different chromosomes; 2) the fusion strains retain the same replication and segregation programs with individual active replication origins and separate, essential partitioning systems; 3) the replication arms on either side of each origin have very similar sizes; and 4) the XerCD/*dif* system is required for the survival of these fusion strains.

The chromosomal fusion sites of the two natural isolates are different from those in the laboratory C58 fusion. The C58 fusion sites were two ∼1.5 kb sequences that differed by only 11 nucleotides (12) (**Fig. 3A**). In contrast, the predicted fusion sequence of CFBP_2407 and CFBP_2642 were identical ribosomal RNA (*rrn*) loci (**Fig. 3B**). There are four nearly identical copies of the *rrn* locus in the genomes of most agrobacteria. In most strains, including 47-2, these are present in two copies on each chromosome, *rrn*1 and *rrn*2 on Ch1 and *rrn3* and *rrn4* on Ch2 (**Fig. S3**). In the two fusion strains we are reporting here, recombination occurred between *rrn*1 on Ch1 and *rrn*3 on Ch2, which resulted in relatively balanced replichores (**Fig. 3B**). Similarly, the C58 fusion chromosome we reported earlier also has roughly balanced replichores (**Fig. 3A**). Since through evolution, replichores are typically balanced in size to maintain correct replication timing (14) and imbalanced replication forks can result in severe fitness effects (15), we speculate that the balanced replichores might be one important reason for the viability of these three fusion strains.

The presence of an ICE immediately adjacent to the fusion site raises questions about its potential role in the fusion process. Mobile genetic elements can mobilize genes important for many phenotypes, however the act of integration can also play a role in microbial evolution (16). Mobile genetic elements such as ICEs have been implicated in facilitating large conformational changes in bacterial chromosomes (17, 18). ICEs target specific sequences, called attachment (*att*) sites, in the chromosome for integration. The predicted *att* site targeted by this ICE is located within the identical repeat region and in multiple copies in the fusion strain and 47-2 genomes. It is plausible that the ICE recombinase may have induced the fusion event by promoting recombination between a Ch1 and Ch2 copy of the *att* site during integration. Future work will investigate whether integration or excision of this ICE from the chromosome can induce fusion or splitting of chromosomes.

Our findings raise important questions about the maintenance and genomic malleability of multipartite genomes. The fusion and splitting of replicons are drivers for genome evolution and speciation. The products of these events can be detrimental to chromosome segregation and can result in extinction of certain lineages from the population. However, it can also be beneficial in some cases. In *V. cholerae*, chromosome fusion was observed when the origin or centromere of one chromosome is mutated; chromosome fusion allowed Ch2 to piggyback on Ch1 so that essential genes were inherited (19). In humans, Robertsonian chromosome translocations are the fusions of two chromosomes which result in viable individuals, but the fused chromosome also has only one functional centromere (20). Here in *A. tumefaciens*, chromosome fusion has been observed in both lab strains and environmental isolates. Importantly, these strains require two origins and two centromeres. The balanced replication arms and a sequence-specific resolution system are the key to the survival of these fusion strains. Our results support the notion that multipartite genomes, and particularly replicon fusions, represent a form of genomic plasticity that can facilitate adaptation but elevates the essentiality of the otherwise non-essential systems like XerCD recombinases to maintain stability. The examination of natural populations of bacteria has now revealed additional examples of chromosome fusions, expanding our understanding of what facilitates and constrains this process. Further exploration for natural examples in more bacterial taxa with multipartite genomes will provide greater insights into the mechanisms controlling genetic diversification and genome evolution.

## Materials and Methods

### Bacteria growth

*A. tumefaciens* strains were grown at 30°C with aeration. For Hi-C and marker frequency analysis, cells were grown in defined minimal medium (ATGN) (21). Single colonies were isolated on ATGN plates and inoculated into 5 ml of ATGN medium and incubated overnight in a tube roller. In the next morning, the cultures were diluted into 30 mL ATGN liquid with a starting OD_600_ of 0.15. Cultures were grown in a shaking water bath for 6 hours to reach an OD_600_ of 0.5 – 0.6 before harvest.

### Nanopore sequencing and analysis

The Wizard Genomic DNA kit (Promega, Madison, WI) was used to extract DNA from overnight cultures of strains 47-2, CFBP_2407, and CFBP_2642. The Rapid Barcoding Kit 96 V14 (SQK-RBK114.96; Oxford Nanopore Technologies, Oxford, UK) was used to prepare multiplexed long read sequencing libraries for these strains. Libraries were sequenced on PromethION flow cells on an Oxford Nanopore P2 solo sequencer with SUP basecalling. Unicycler v0.5.0 with the parameter “--mode bold” was used to generate hybrid assemblies for strains CFBP_2407 and CFBP_2642 (22). For strain 47-2, Canu v2.2 with the parameters “useGrid=false genomesize=5.6m” was used to assemble the genome from long-read sequences (23). Polypolish v0.5.0 with the default parameters was used to polish the long-read assembly with Illumina short reads (24).

Publicly available *Agrobacterium* genomes were downloaded from NCBI on July 19, 2023. Beav v1.0.0 with parameters “--agrobacterium--tiger_blast_database” and a blast database of the public genome dataset was used to annotate genomes (25). FastANI v1.34 with the default parameters was used to calculate pairwise average nucleotide identity (ANI) between all strains (26). AutoMLSA2 v.0.9.0 with the default parameters was used to generate a multi-locus sequence analysis (MLSA) phylogeny of all publicly available *Agrobacterium* strains and the two fusion strains (27). Twenty-three translated reference gene sequences from strain C58 were used as queries for the MLSA analysis. Genes used were *acnA*, *aroB*, Atu0781, Atu1564, Atu2640, *cgtA*, *coxC*, *dnaK*, *glyS*, *ham1*, *hemF*, *hemN*, *hom*, *leuS*, *lysC*, *murC*, *plsC*, *prfC*, *rplB*, *rpoB*, *rpoC*, *secA* and *truA* (28). PIRATE v1.0.5 with the default parameters was used to cluster genes from 6 strains belonging to the MLSA tree clade containing CFBP_2407 and CFBP_2642 into orthologous groups (29). To prepare input files for PIRATE, the agat v1.4.0 script agat_convert_sp_gxf2gxf.pl with the “--gff_version_output 3” parameter was used to repair GFF3 files prior to ortholog clustering (30). MAFFT v7.525 with the default parameters was used to align sequences of 4,192 conserved genes from all genomes (31). The catfasta2phyml v.1.2.0 tool with parameters “-f--concatenate” was used to concatenate gene alignments (32). IQ-TREE2 v2.3.1 with the parameters “-bb 1000-alrt 1000” was used to infer a maximum likelihood phylogeny from the core genome alignment (33). A custom Python script was used to generate synteny plots (https://github.com/acarafat/PySyntenyViz).

### Chromosome Confirmation Capture (Hi-C)

The Hi-C procedure performed here was as previously described (12, 34). In brief, we grew *A. tumefaciens* strains to exponential phase (∼0.6 OD_600_) in ATGN broth at 30°C (21) before cells were cross-linked with 3% formaldehyde at room temperature for 30 min. Crosslinking was quenched with 125 mM glycine for 5 min and spun down for pellets normalized to 1.2 OD_600_ units. Pellets were lysed using Ready-lyse lysozyme (Epicentre, R1802M) then treated with 0.5% SDS. Solubilized chromatin was digested with HindIII for 2 hours at 37°C. Digested DNA ends were filled in with Klenow DNA polymerase and Biotin-14-dATP, dGTP, dCTP, dTTP. Biotinylated products were ligated with T4 DNA ligase at 16°C for about 20 hr. The DNA products were subsequently reverse crosslinked at 65°C for about 20 hr in the presence of EDTA, proteinase K, and 0.5% SDS. The reactions were extracted twice using phenol/chloroform/isoamylalcohol (25:24:1) (PCI), precipitated with ethanol, and resuspended in 20 µl of 0.1XTE buffer. Biotin from non-ligated ends were removed using T4 DNA polymerase (4 hr at 20°C) followed by extraction with PCI. The DNA was then sonicated for 12 min with 20% amplitude using a Qsonica Q800R2 water bath sonicator. Sheared DNA was library prepped using NEBNext UltraII kit (E7645) and biotinylated DNA fragments were purified using 10 µl streptavidin beads. DNA-bound beads were used for PCR in a 50 µl reaction for 14 cycles. PCR products were purified using Ampure beads (Beckman, A63881) and sequenced at the Indiana University Center for Genomics and Bioinformatics using NextSeq2000. Paired-end sequencing reads were mapped to the combined hybrid assemblies for strains CFBP_2407, CFBP_2642 and 47-2 described in “Nanopore sequencing and analysis” methods. Reads in which both reads uniquely aligned to the genome were processed and sorted by HindIII restriction fragments and binned into 10-kb bins using HiC-Pro pipeline (35). Analyses and visualizations were done using R. 47-2 genome was rearranged for *ori1* at the center of Ch1 with the following rearrangement: Ch1 1040.763 - 2893.626 kb, 0.001 kb - 1040.762 kb.

### Whole genome sequence (WGS) and marker frequency analysis (MFA)

Cells were grown to exponential phase (∼0.6 OD_600_; 5×10^9^ CFU/mL) in ATGN broth at 30°C (21) in which 5 mL was collected and cells pelleted by centrifugation at 5000 xg for 10 min. Genomic DNA was extracted using Qiagen DNeasy Kit (69504), sonicated using a Qsonica Q800R2 water bath sonicator at 20% continuous amplitude for 6 min.

Sheared gDNA was library prepared using the NEBNext UltraII kit (E7645), and sequenced using Illumina NextSeq2000. The reads were mapped to the combined hybrid assemblies for strains CFBP_2407, CFBP_2642 and 47-2 using CLC Genomics Workbench (CLC Bio, QIAGEN). The mapped reads were normalized by the total number of reads. Relative copy numbers were calculated by dividing normalized reads with the averaged total number of reads at the terminus of Ch1 or at the left terminus of the fused chromosome. Plotting and analysis were performed using R.

### Transposon insertion sequencing (Tn-seq)

Tn-seq was performed as previously described (12). In brief, the *Mariner* transposon-based plasmid pTND2823 (gift from Triana Dalia and Ankur Dalia at Indiana University) was transformed into diaminopimelic acid auxotroph *MFDpir E. coli* (36) used to conjugate with *A. tumefaciens* strains. *A. tumefaciens* conjugants were plates on 10 large plates (150 mm diameter, VWR 25384-326) with LB supplemented with 300 mg/ml kanamycin aiming for 1×10^6^ kanamycin-resistant colony forming units. The plates were incubated at 30°C for two days before each plate was scraped for all colonies and combined into a single pool. 5 OD_600_ unit of cells from the pool was used for genomic DNA (gDNA) isolation using QIAGEN DNeasy blood & tissue kit (69504). 3 µg of gDNA was digested with MmeI (NEB R0637S) for 90 min, and treated with Calf Intestinal Alkaline Phosphatase (Quick CIP, NEB M0525L) for 60 min at 37°C. The DNA was extracted using Phenol-Chloroform, precipitated using ethanol and resuspended in 15 µl ddH2O. The digested end was ligated to an annealed adapter49 using T4 DNA ligase and incubated at 16°C for about 16 hr. Adapter-ligated DNA was amplified with the primers complementary to the adapter and the transposon inverted repeat sequence.

The PCR product was gel purified using a Monarch DNA Gel Extraction Kit (NEB T1020S) and sequenced at the IU Center for Genomics and Bioinformatics using NextSeq2000. Sequencing reads were mapped to the combined hybrid assemblies for strains CFBP_2407, CFBP_2642 and 47-2 and analyzed using a previously described approach (37, 38). The results were visualized using Artemis (https://www.sanger.ac.uk/tool/artemis/).

## Acknowledgements

We thank IU CGB for next-generation sequencing, Ankur Dalia for the transposon plasmid, and Doron Teper and Shulamit Manulis for sharing strain 47-2. This work was supported by the NIH (R01GM141242, R01GM143182, and R01AI172822 to X.W.), NSF GEMS Biology Integration Institute (2022049 to X.W. and C.F.), and startup funds from Oregon State University (to A.R. and A.J.W.),

## Supplementary Information for

**Fig. S1.**
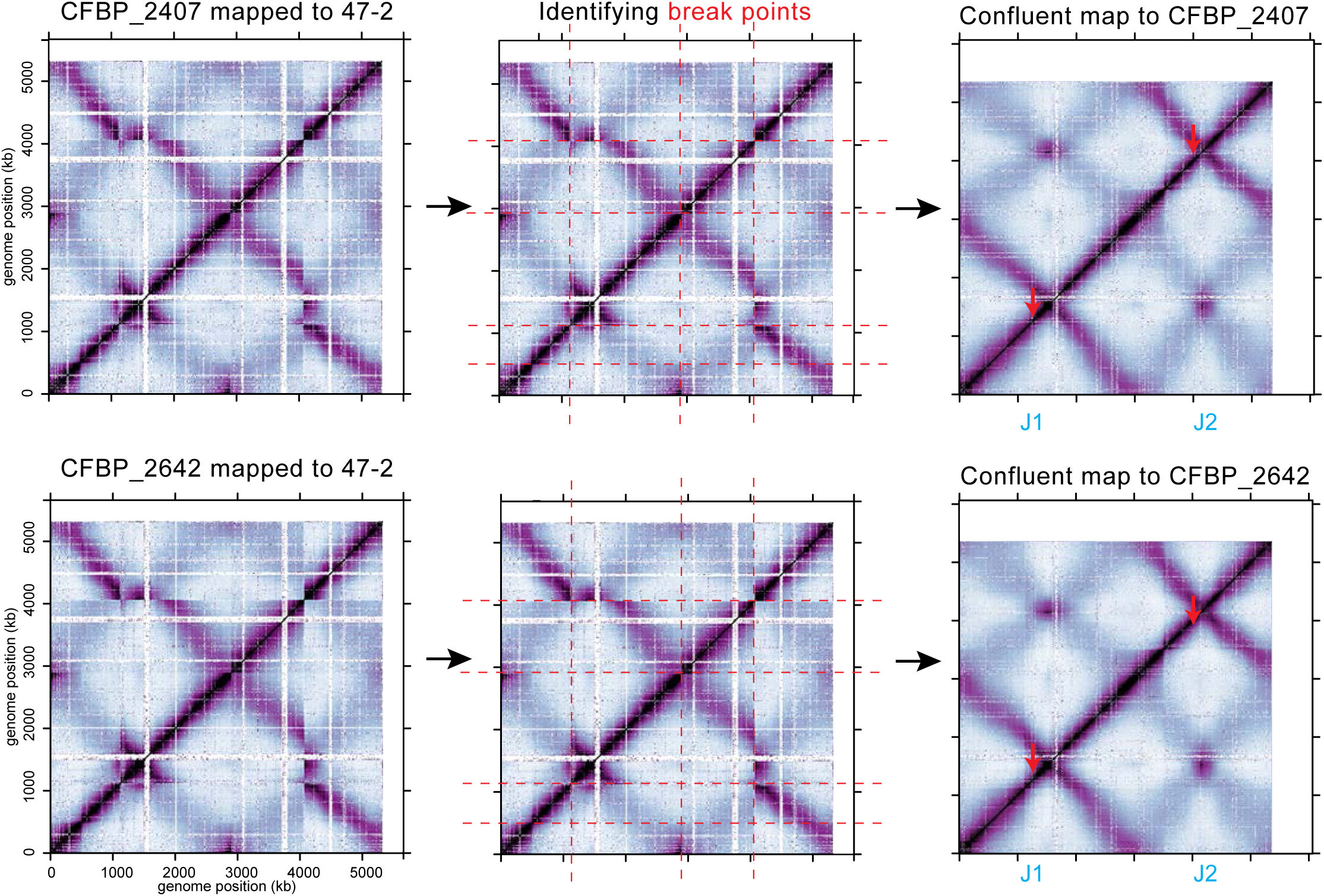
Identifying the junction of chromosome fusion. Left: Hi-C reads from fusion isolates (CFBP_2407 and CFBP_2642) were mapped to the genome of the binary 47-2 strain. White spaces indicate unmapped regions on the 47-2 genome. Middle: The breakpoints of the Hi-C maps were highlighted with red dashed lines. The maps were cut and reassembled to best match the confluent maps generated using sequenced genomes (Right). Based on this approach, we identified genetic loci of the probable fusion junctions (red arrows).

**Fig. S2.**
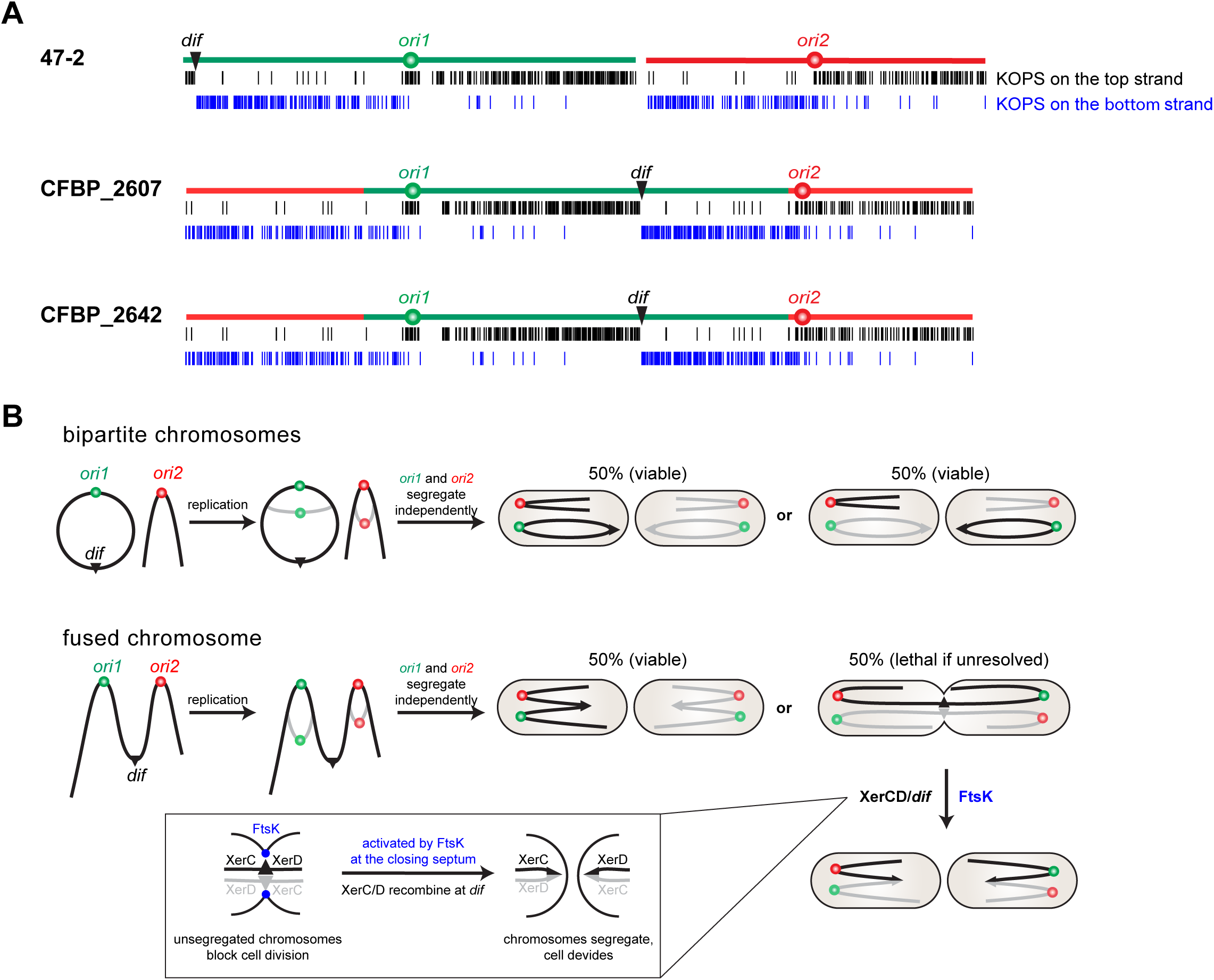
Fusion chromosomes require the XerCD*/dif* system for survival. **(A)** Distribution of the FtsK orienting polar sequences (KOPS) (GGGNAGGG) on the chromosomes of the indicated strains. Black and blue bars indicate KOPS sequences on the top and bottom DNA strand, respectively. The *dif* site is illustrated by a black triangle. **(B)** Schematics models adapted from (1). After replication initiation, bipartite chromosomes separate independently and randomly into daughter cells. Both combinations will result in viable cells. For the fused chromosome, random segregation of *ori1* and *ori2* will generate 50% of cells with intact fusion chromosomes in each daughter cell, resulting in viable progeny. However, in 50% of the cases in every generation, the copies of *ori1* and *ori2* that belong to the same DNA molecule translocate to opposite daughter cells. If the situation is unresolved, the chromosomes will block cell division and cause cell death. Stimulated by FtsK at the closing septum, XerCD will recombine at the *dif* sites to resolve the unsegregated chromosomes.

**Fig. S3.**
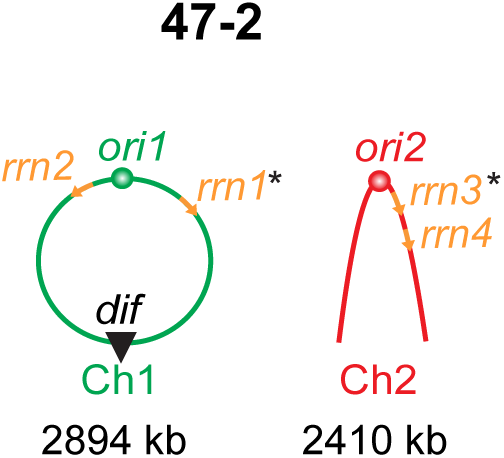
Locations of *rrn* operons on the chromosomes of strain 47-2. Asterisks mark the sites of chromosome fusion in CFBP_2407 and CFBP_2642.

**Table S1.** Bacterial strains used in this study.

| Strain | Reference | Figure |
| --- | --- | --- |
| *A. tumefaciens* 47-2 (BV1 biovar; env. isolated binary chromosome) | This Study | 1, 2A, 3BC, 4, S1, S2 |
| *A. tumefaciens* CFBP_2407 (BV1 biovar; env. isolate fused chromosome) | This Study | 1, 2B, 3BD, S1, S2 |
| *A. tumefaciens* CFBP_2642 (BV1 biovar; env. isolate fused chromosome) | This Study | 1, 2C, 3BD, S1, S2 |

**Table S2.** Nanopore sequencing samples generated in this study.

| Sample name | Figure | Reference |
| --- | --- | --- |
| 47-2 | 1AB | This Study |
| CFBP_2407 | 1AB | This Study |
| CFBP_2642 | 1AB | This Study |

**Table S3.**
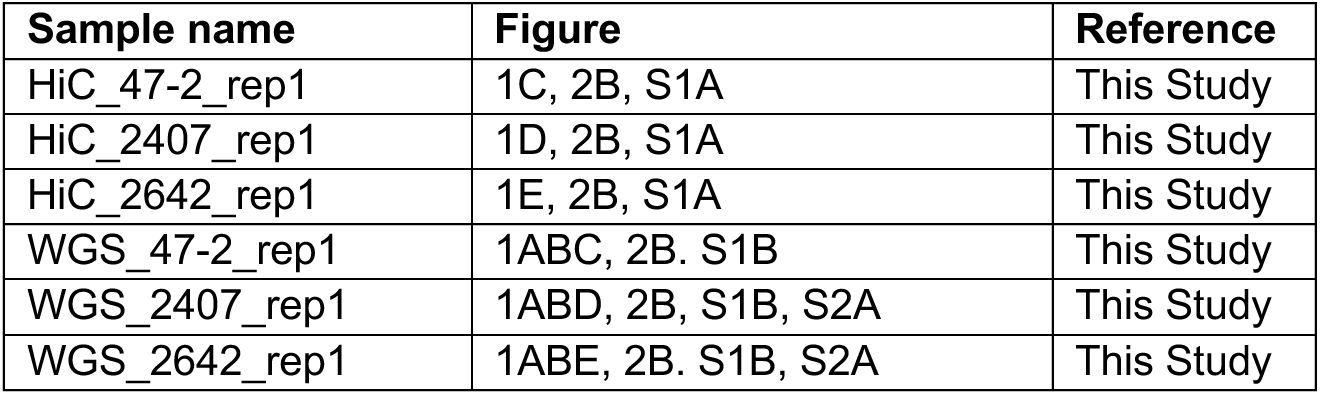
Hi-C and WGS samples generated in this study.

| Sample name | Figure | Reference |
| --- | --- | --- |
| HiC_47-2_rep1 | 1C, 2B, S1A | This Study |
| HiC_2407_rep1 | 1D, 2B, S1A | This Study |
| HiC_2642_rep1 | 1E, 2B, S1A | This Study |
| WGS_47-2_rep1 | 1ABC, 2B, S1B | This Study |
| WGS_2407_rep1 | 1ABD, 2B, S1B, S2A | This Study |
| WGS_2642_rep1 | 1ABE, 2B, S1B, S2A | This Study |

## Notes

### Competing Interest Statement

The authors have declared no competing interest.

